# Comparative study of contact repulsion in control and mutant macrophages using a novel interaction detection

**DOI:** 10.1101/2020.03.31.018267

**Authors:** JA Solís-Lemus, BJ Sánchez-Sánchez, S Marcotti, M Burki, B Stramer, CC Reyes-Aldasoro

## Abstract

This paper compares the contact-repulsion movement of mutant and wild-type macrophages using a novel interaction detection mechanism. The migrating macrophages are observed in *Drosophila* embryos. The study is carried out by a framework called `macrosight`, which analyses the movement and interaction of migrating macrophages. The framework incorporates a segmentation and tracking algorithm into analysing motion characteristics of cells after contact. In this particular study, the interactions between cells is characterised in the case of control embryos and Shot3 *mutants*, where the cells have been altered to suppress a specific protein, looking to understand what drives the movement. Statistical significance between control and mutant cells was found when comparing the direction of motion after contact in specific conditions. Such discoveries provide insights for future developments in combining biological experiments to computational analysis. Cell Segmentation, Cell Tracking, Macrophages, Cell Shape, Contact Analysis

## 1 INTRODUCTION

Cellular migration is a process of of great importance in many biological processes, for instance endothelial migration [1], infection [2], angiogenesis [3], tissue regeneration [4], inflammation [5, 6, 7, 8, 9], cellular invasion by parasites [10], metabolic changes in macrophages due to hypoxia [11], cancer invasion and metastasis [12, 13, 14], embryogenesis [15, 16], healing of wounds [17, 18] and in the context of this work, the development of the immune system [19, 20, 21]. Together with neutrophils, macrophages are important cells of the immune system [22, 6]. One of the functions of macrophages is to filter foreign particles when settled in lymphoid tissues and the liver [19]. Macrophages have an important role in homeostasis [23], which is the tendency of a system to keep the equilibrium of physiological processes. In these cases, the role of macrophages ranges from tissue repair through to immune responses to pathogens [24]. The presence of macrophages can be a signal of problems. Excessive migration can be related to autoimmune diseases and cancer [25] as the macrophages can be related to fighting altered cells.

*Drosophila melanogaster*, also known as the common fruit fly, has been widely studied as a model organism [26, 27, 28]. Although in evolutionary terms the fly is very far from vertebrates, it shares many developmental and cellular processes with other organisms, including humans [29]. Thus, investigations with Drosophila have led to insights about the role of macrophages and how they integrate migratory movement with external cues [20]. For instance, previously unknown movements of cytoskeletal structures in macrophages were discovered, specifically, the cell-cell contacts appeared to alter migration [30, 31]. In **(author?)** [31], contact inhibition of locomotion was described in these cells, showing that these cell-cell interactions are needed for functional migration.

**Tracking of cells** comprises the identification of the cells from background and then linking between previously detected cells in one time frame to the same cells in subsequent frames. In this work, tracking will be defined as a function of **segmentation**, that is, the correct identification of each cell from the background, and probably more important, and from the other cells. Both segmentation and tracking of cells have been widely studied with many imaging modalities [32, 33, 34, 35]. Cell tracking when cells are observed with phase contrast microscopy have been presented in [32, 33], showing quantitative analysis of cell dynamics *in vitro*. In [34, 35], several tracking methodologies were evaluated with a series migratory cells under different conditions. The methodologies were compared, not only in their ability to track the cells that were segmented, but also to identify event like mitosis. Other cellular events, e.g. interactions between cells, are also of huge importance as these may be related to communication between cells or cell signalling. To study these events, a more thorough study of a tracks’ features is necessary.

**Movement analysis**, in this work will be defined as the analysis of features derived from tracks, and will be performed to examine specific research questions related to certain phenomena to be studied. For instance, in [36], tracks are classified depending on certain features, e.g. curvature and speed. In a related work, a movement pattern analysis provided insights about a toxicological environment assessment with *Artemia Franciscana* swimming in chambers with sub-lethal doses [37]. In that experiment, the tracks produced by the movement of these marine crustaceans were examined for specific patterns of migration (circular motions) which were related to the levels of toxicity. Contributions regarding the specific data analysed in this work have been varied. Segmentation of macrophages in single frames was presented in [38], showcasing the complex interactions which manifest as overlapping (*clumps*). In [39], the relationship between contiguous frames was incorporated to the segmentation of single cells, allowing for a controlled measurement of shape parameters between overlapping events. Finally, `macrosight`, a software framework to analyse the movement and the shape variation of fluorescently labelled macrophages was presented in [40], where overlapped clumps are considered moments of assumed interaction between the cells and thus the movement before and after contact was analysed.

In the present study, macrophages from control embryos were compared to *Shot3* embryos, referred to as *mutants. Shot3* is a mutation which affects the cytoskeletal crosslinking [41, 42]. The `macrosight` software is used to search for an underlying difference in the movement between controls and mutant time sequences. The main contribution consists of the use of a software framework to provide robust, quantitative measurements of the same object in different conditions. It is worth noting that the two main hypotheses of `macrosight` are (i) that cell-cell contact accounts for an interaction between cells, and (ii) as a result of an interaction, one or both cells involved in the interaction will noticeably change the direction before and after contact. Furthermore, Figure 1 shows a graphical abstract of the main contribution of the comparison between control and mutant experiments.

**Figure 1:**
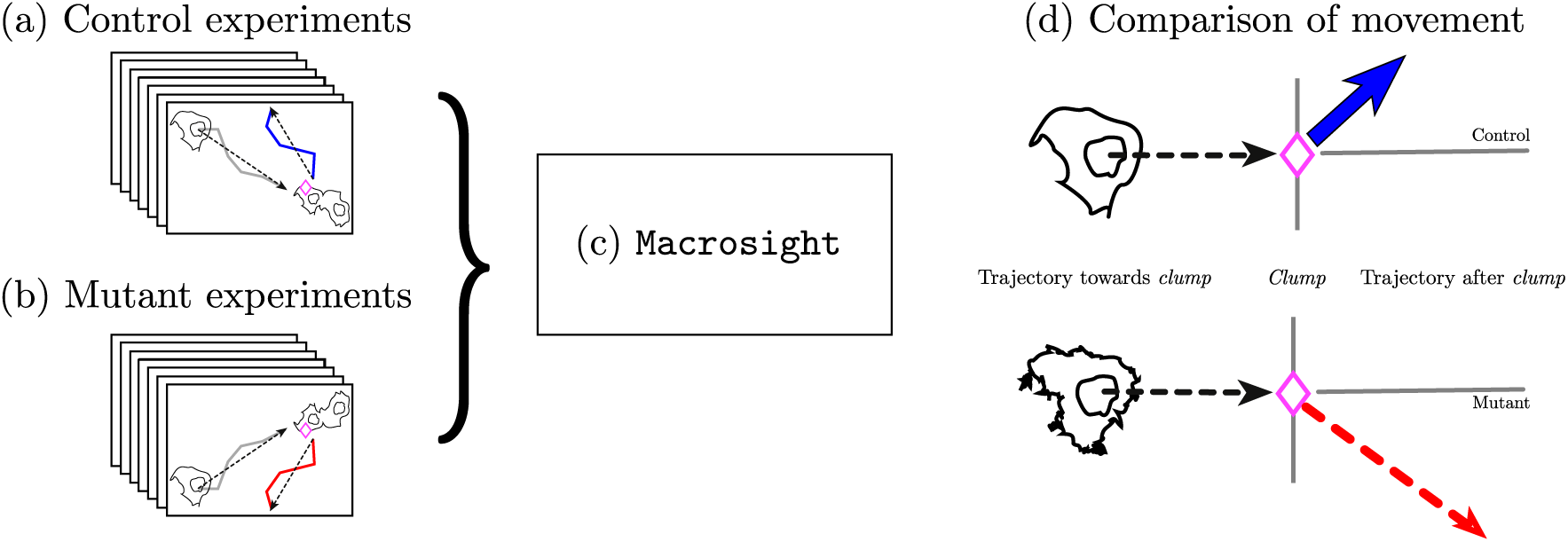
Illustration of the main hypothesis in this work. Differet movement patterns from control to mutant experiments are expected as a result of the movment analysis performed. The diagram shows the two different types of cells: controls (a) and mutants (b) being procesed with `macrosight` [40] (c). The output (d) consists of measurements of the cell’s trajectories and the changes in direction upon interactions represented by the different types of line and colours in the diagram.

A preliminary version of this work was presented at the 23rd Medical Image Understanding and Analysis (MIUA) [43]. The algorithms have been extended and several new experiments with new data are presented. Thus, this work now describes the following topics, not included previously: (i) a more thorough explanation on the interactions of macrophages and a stronger description of the methodology; (ii) new representation of distribution of angles, allowing a much better interpretation of the results; and (iii) a more thorough literature review of the problem.

## 2 MATERIALS

In this work, a total of 14 time sequences of macrophages in *Drosophila* embryos were analysed. The macrophages were observed with fluorescence microscopy following the protocol described in [30, 31]. The nuclei were labelled in red and the microtubules were labelled in green. Each image of a time-lapse sequence was acquired every ten seconds, and the lateral dimensions of the pixel were 0.21*µm*. The dimensions of the images of the fourteen experiments were (*n*_*w*_, *n*_*h*_, *n*_*d*_) *=* (512, 672, 3) [rows, columns, channels].

Two different types of cells were acquired, i.e. wild-type controls and mutant experiments. Four control and ten *Shot3* mutant experiments were analysed. The number of time frames of the control data sets ranged between 137 and 272, whilst for the mutant it was between 135 and 422 frames. Figure shows a comparison with four frames of one control and one mutant.

The overlap of two cells, which are defined as *clumps*, are very important for the study of interactions caused by cell-cell contact, as presented in Fig, 3.

**Figure 2:**
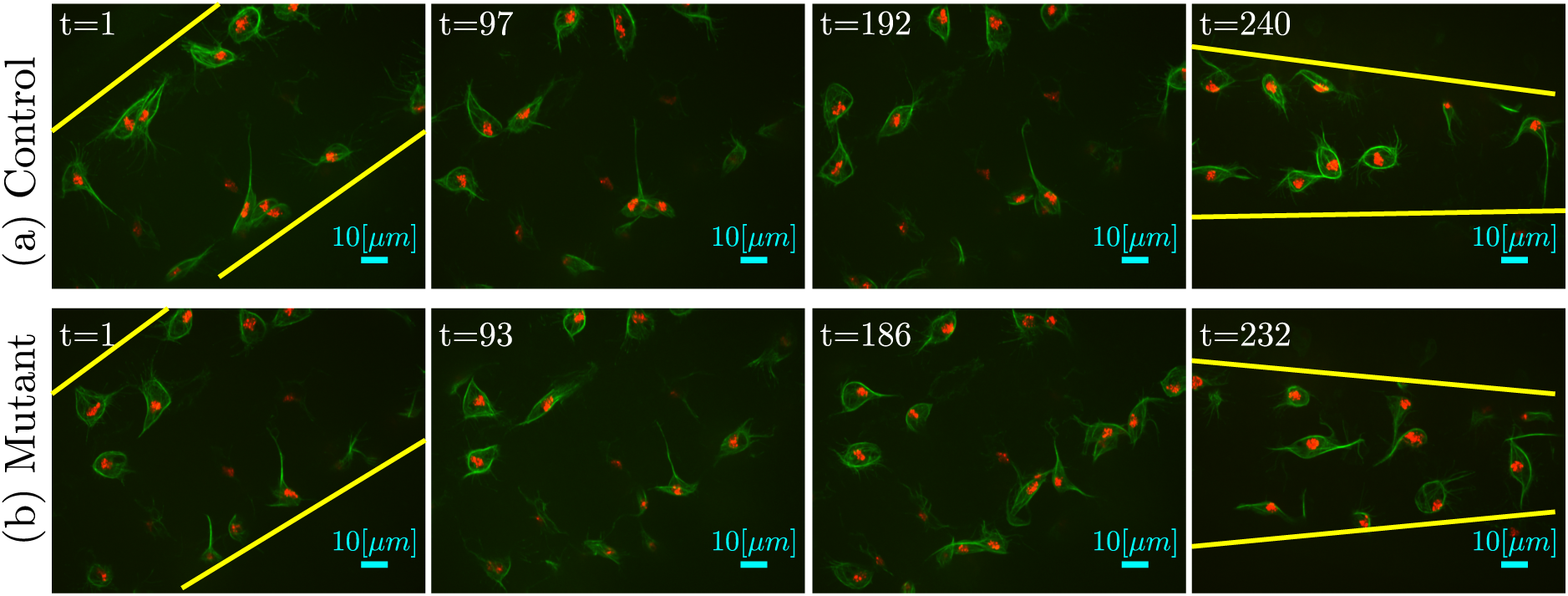
Comparison between four frames of (a) control against four frames of a (b) mutant data set. These data sets were selected as they had a similar number of frames and thus a similar spacing between the frames in both cases could be shown (≈95). Yellow lines have been manually inserted to the initial and final frames on both experiments to illustrate the apparent change of focus of the microscope as time evolves.

**Figure 3:**
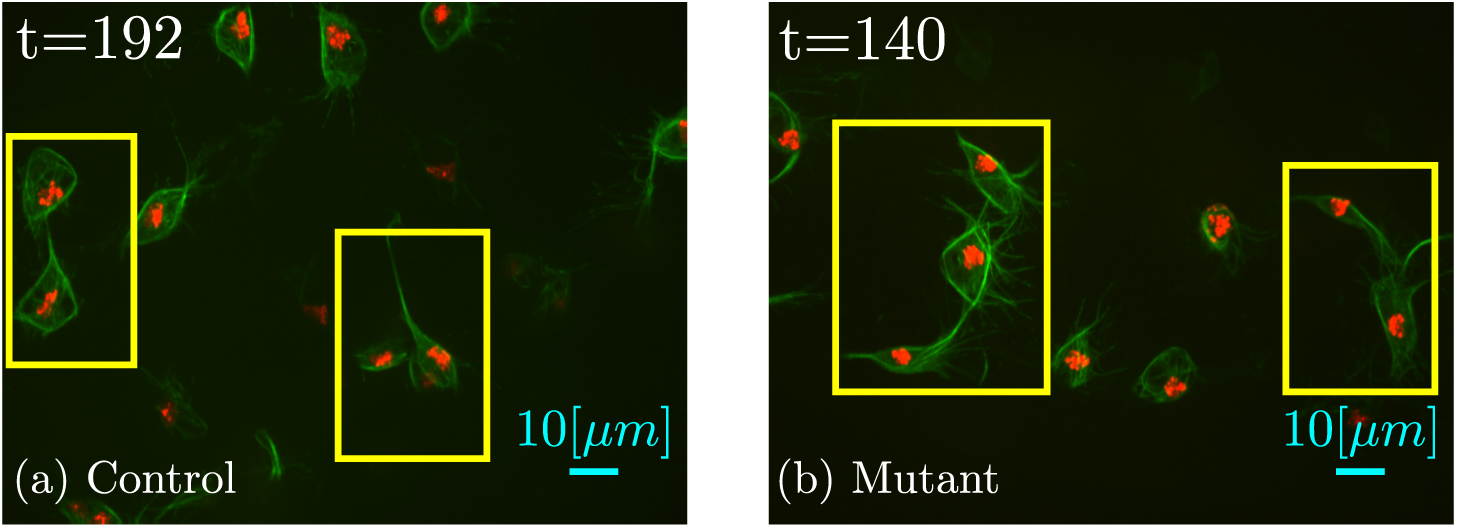
Illustration of a series of *clumps* in (a) control and (b) mutant experiments. Both data sets present overlapping events, i.e. *clumps* which are highlighted with yellow boxes. It should be noticed that although the microtubules are overlapping, the nuclei are still separated.

Under certain circumstances, cells have been shown to align their microtubules and change drastically the orientation of movement [31]. The contact observed in certain *clumps* suggest a change of direction of the migration patterns of those cells involved in the contact. This type of interaction was analysed previously in [40], where cell-cell contact was shown to be influence the movement of cells.

## 3 METHODS

`Macrosight` [40] is a framework for the analysis of moving macrophages capable of segmenting the two layers of fluorescence in the dataset presented previously, and apply the keyhole tracking algorithm inside the PhagoSight framework [44] on the centroids of the segmented nuclei. Figure 4 shows an illustration of the flow of information in `macrosight`. Each track generated *𝒯*_*r*_ contains information on the (i) position **x**_*t*_ at a given time frame *t*, (ii) track identifier *r*, (iii) velocity *v*_*t*_, and whether the cell is part of a clump.

**Figure 4:**
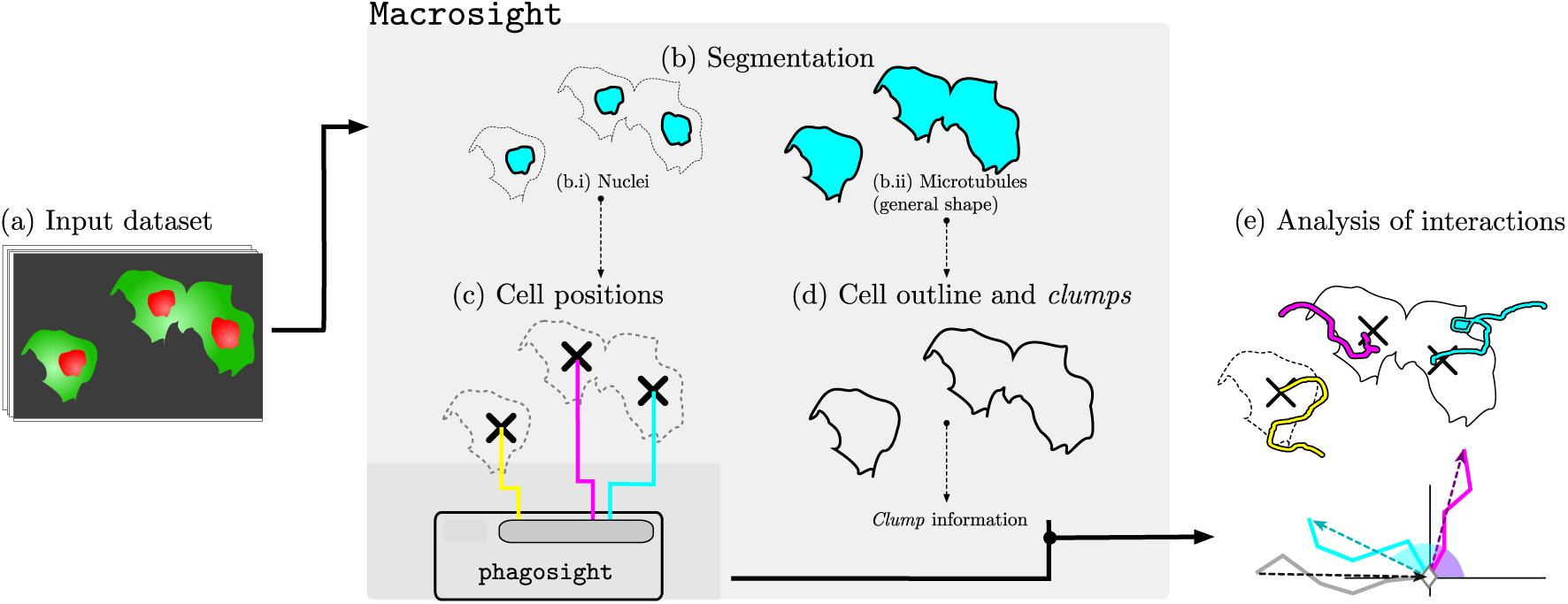
Illustration of the `macrosight` framework parts used in this work. (a) Illustration of a sequence of images with cells with red nuclei and green microtubules. The two fluorescent channels are segmented in (b) based on a hysteresis threshold where the levels are selected by the **(author?)** [45] algorithm. The segmentation of the red channel (b.i) provides the cell positions necessary to produce (c) the tracks of the cells using the keyhole tracking algorithm [44] (represented in cyan, magenta and yellow). Finally, the tracks’ information is combined with the clump information (d) from the segmented green channel (b.ii) to allow analysis of movement based on contact events (e), producing the change of direction chart per cells in clump.

Each *clump* can be uniquely identified through an individual code *c*(*r, q*), where *r* > *q* indicates that at a certain time frame *t*, tracks 𝒯_*r*_ and 𝒯_*q*_ belong in the same clump. This allows for each interaction to be analysed. Several tracks can join together into a single clump, thus the *clump* codes evolve. Figure 5 illustrates the evolution of a given track *𝒯*_2_ and its involvement in two different clumps as a cartoon.

**Figure 5:**
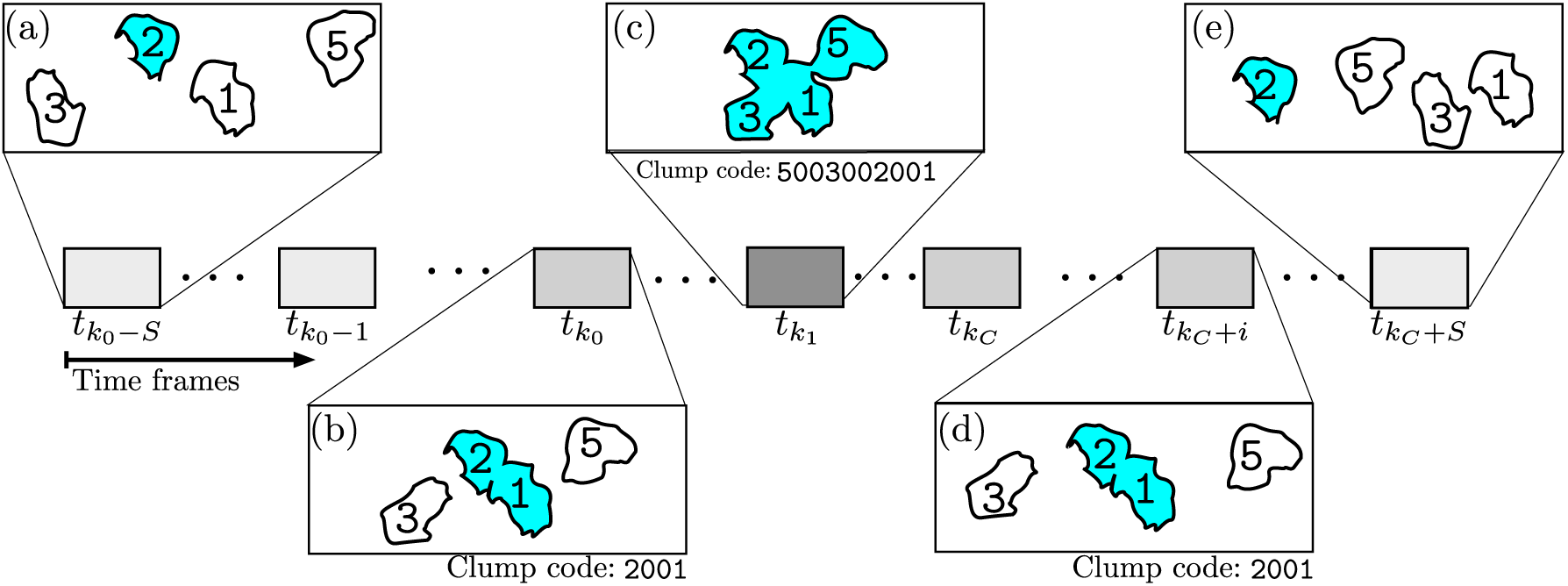
Illustration of clump codes to the different time frames for a particular track *𝒯*_2_. The horizontal axis represents the time, and the detail of five frames is presented to illustrate the evolution of track *𝒯*_2_ as it interacts with other cells. In (a) and (e), track *𝒯*_2_ is not in contact with any other cell, thus no clump is present. (b) Represents the moment when *𝒯*_2_ and *𝒯*_1_ start interacting in clump `2001`. Following in (c), tracks *𝒯*_3_ and *𝒯*_5_ become present in the clump, thus the *clump* code changes to `5003002001`.

To provide the reader with a real representation of the cell movements, Figures 6 and 7 show qualitatively the movement of the cells before and after a contact event. The number of frames the cells appear in a *clump* is relevant to the study of the movement as it acts as a proxy for the time cells were in actual contact (ten seconds per frame). In the examples shown, cells in Figure 6 remain in the clump for 6 frames (one minute), while the cells in Figure 7 are shown in the clump for 18 frames (3 minutes). It is worth noting that a single clump could provide more than one experiment in different time spans, as the two interacting tracks could interact with each other back and forth. An illustration of one interaction is shown in Figure 8.

**Figure 6:**
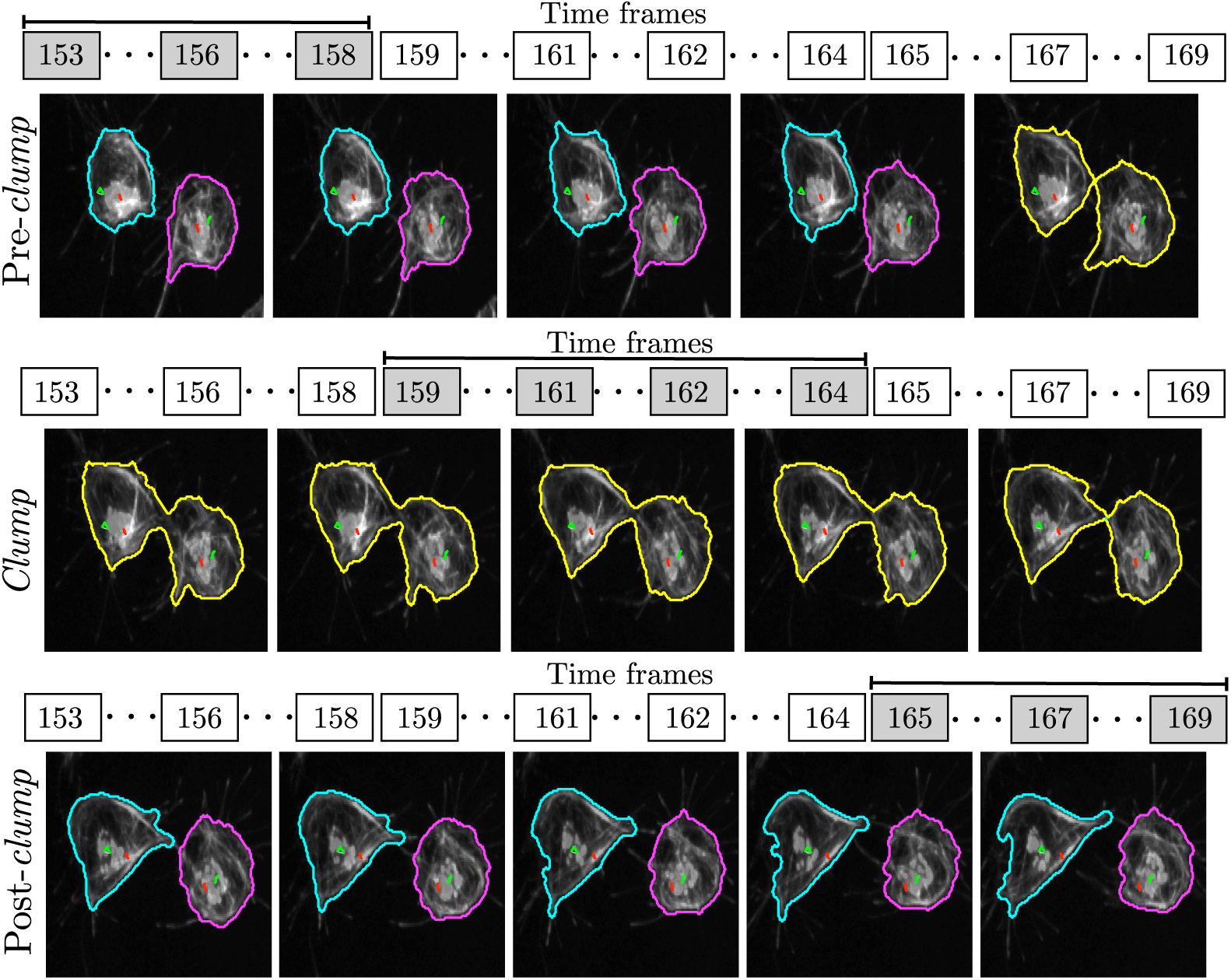
Representation of the migration of two cells as they form a *clump*. The detected clump outlines are presented in yellow line and the individual cells are shown in cyan and magenta. To show the time the duration of the *clump*, the specific time frames are shown, in this case, the clump lasting for 6 frames (60 seconds).

**Figure 7:**
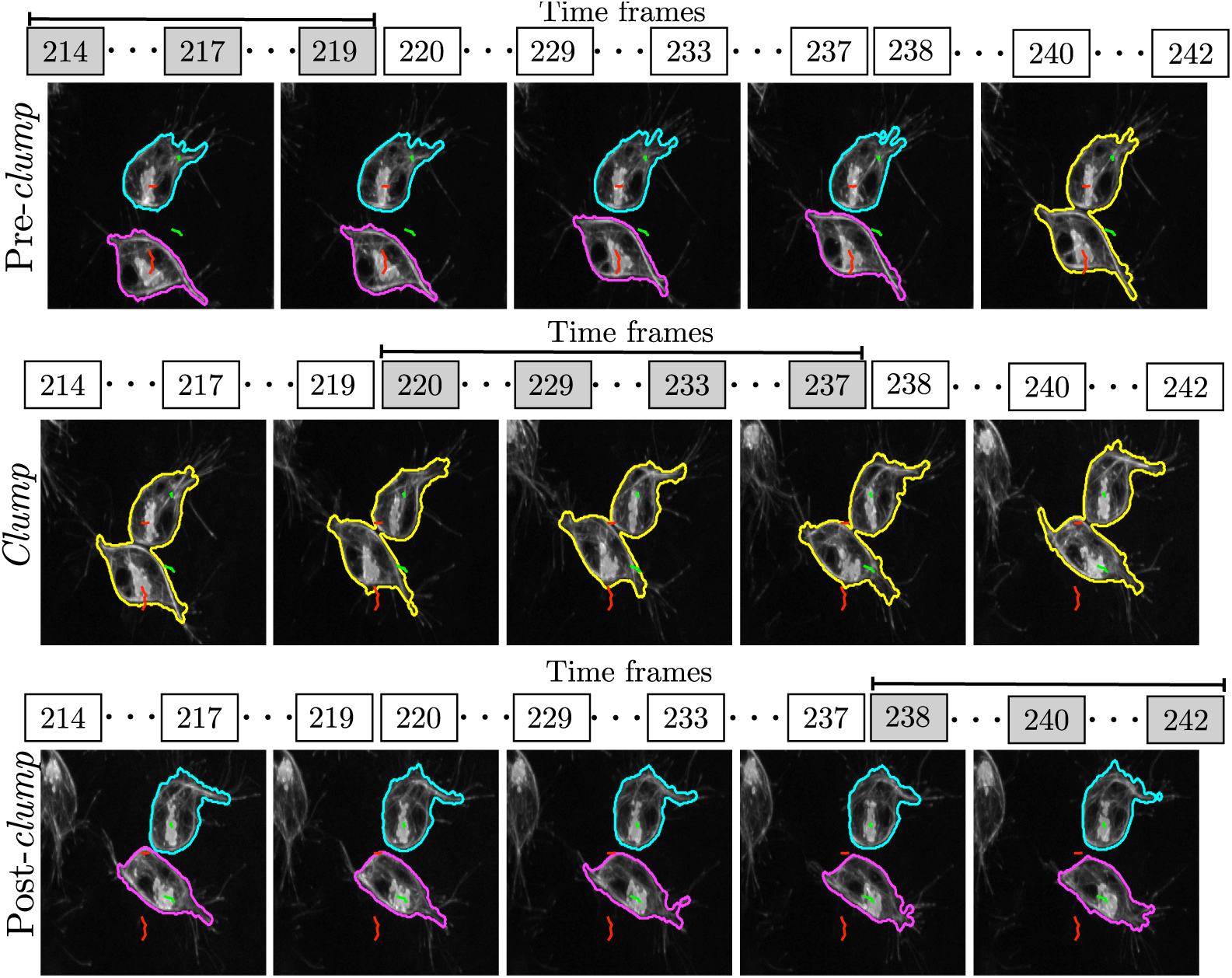
Representation of the migration of two cells as they form a *clump*. The detected clump outlines are presented in yellow line and the individual cells are shown in cyan and magenta. To show the time the duration of the *clump*, the specific time frames are shown, in this case, the clump lasting for 18 frames (180 seconds).

**Figure 8:**
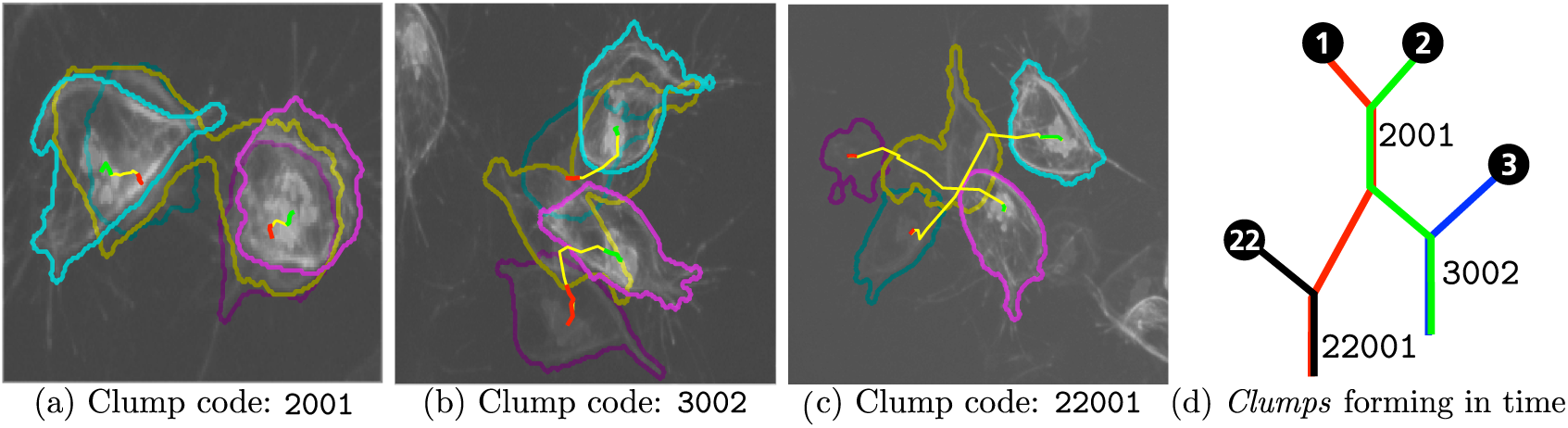
Frames in different interactions overlapped to appreciate cell movement and *clump* formation. (a-c) Three frames are superimposed: the first, middle and final frames in each experiment are shown, with corresponding segmentations and tracks. The full track in each experiment is presented, with changes of colour representing different moments: before (red), during (yellow) and after (green) the *clump*. (d) Is a representation of the same cells forming different *clumps* at different time points.

### 3.1 Analysis of Movements and Interactions

The events of interest in this paper consist of analysing cell-cell contact events of two cells, these will be called *interactions*. The change of direction *θ*_*x*_ ∈ (−*π, π*) is calculated by taking the positions of the tracks *𝒯*_*r*_ and *𝒯*_*q*_ up to *S* frames prior the first contact at time frame 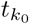, as well as the positions up to *S* frames after the last time frame of contact 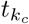. The time in clump 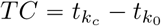 refers to the number of frames the two tracks interact in a given instance of the clump, and it is not taken into consideration for the calculation of angle *θ*_*x*_. A diagram of the calculation of *θ*_*x*_ is provided in Figure 9, where the positions on the image **x=**(*x, y*) get translated and rotated into new frame of reference (*x*′, *y*′).

**Figure 9:**
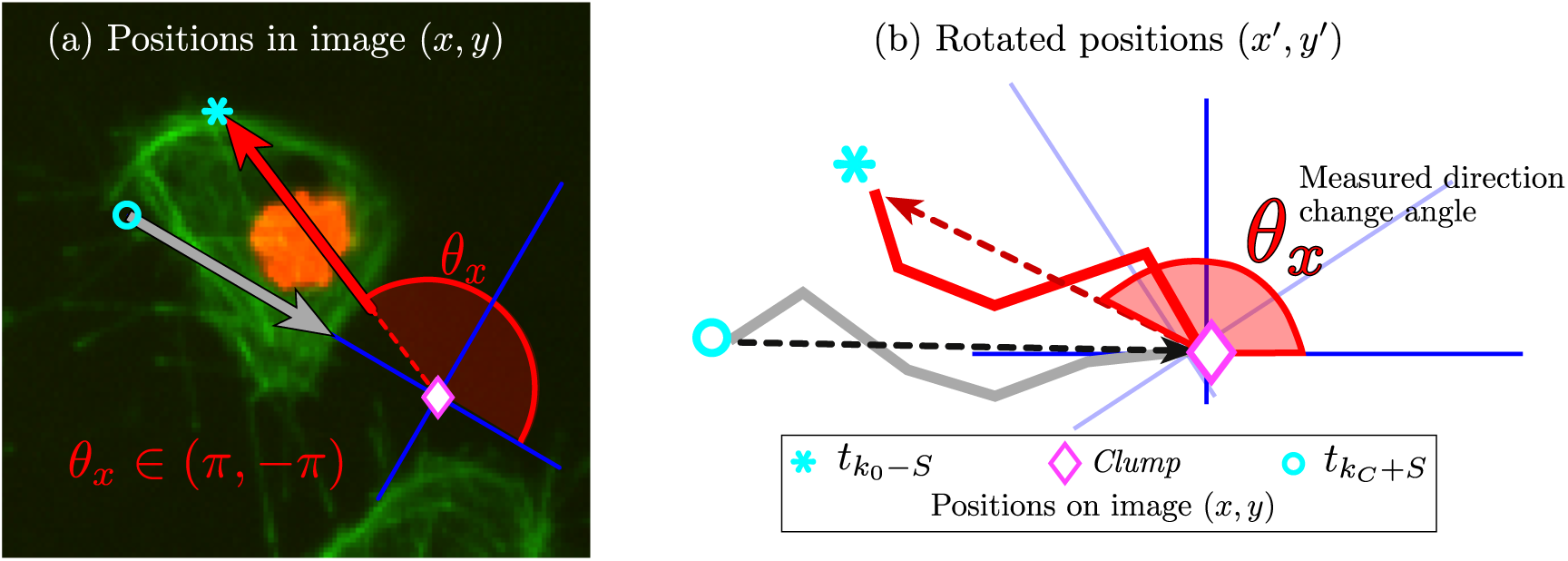
Illustration of direction change (*θ*_*x*_) measurement. Three markers represent different positions of a given track. The markers are (○) represents *S* frames before contact; (◊) represents the starting instant of the clump; and (*) represents the position where the experiment is finalised. Notice the translation and rotation into the new frame of reference (*x*′, *y*′).

### 3.2 Selection of interactions

All available datasets were segmented and tracked. The tracks’ information was searched to find types of *clumps* were considered as interactions, i.e. those which fulfilled the following criteria: (i) **only two cells interacting**; (ii) **full interaction**, where at least one of the cells would enter and exit the *clump*; and (iii) **immediate reaction**, with a value of *S* ranging from 3 to 5, which corresponds to 30 to 50 seconds. The changing direction angles, *θ*_*x*_, for each case were calculated, recording the time in clump *TC* and change of direction.

## 4 RESULTS

After the processes of segmentation, tracking, and selection of suitable interactions, twenty four control and thirty nine mutant interactions were selected for analysis. Table 1 shows the number of interactions per dataset selected, it is worth noting the different numbers of interactions fitting the criteria between datasets.

**Table 1:** Number of suitable interactions per dataset. Notice that not all datasets provided the same number of interactions for the analysis, as one or more of the selecting criteria would not be fulfilled. Also, the mutant datasets 01,02 and 09 did not provide any suitable interactions due to clumps always involving more than two cells.

| Dataset ID | $n$ interactions | Dataset ID | $n$ interactions | Dataset ID | $n$ interaction |
| --- | --- | --- | --- | --- | --- |
| CONTROL01 | 14 | MUTANT03 | 10 | MUTANT07 | 3 |
| CONTROL02 | 4 | MUTANT04 | 2 | MUTANT08 | 2 |
| CONTROL04 | 4 | MUTANT05 | 2 | MUTANT10 | 4 |
| CONTROL05 | 2 | MUTANT06 | 9 | MUTANT11 | 7 |
| TOTAL | 24 | TOTAL | 39 |  |  |

The number of interactions per dataset averaged 6 ± 5.41 for controls and 4.87 ± 3.31 for mutants. The resulting tracks representing changes of direction are shown in figure 10 for (a) control and (b) mutant. Differences can be observed in the displacement of the cells towards and from the centre, in the horizontal direction *x*′.

**Figure 10:**
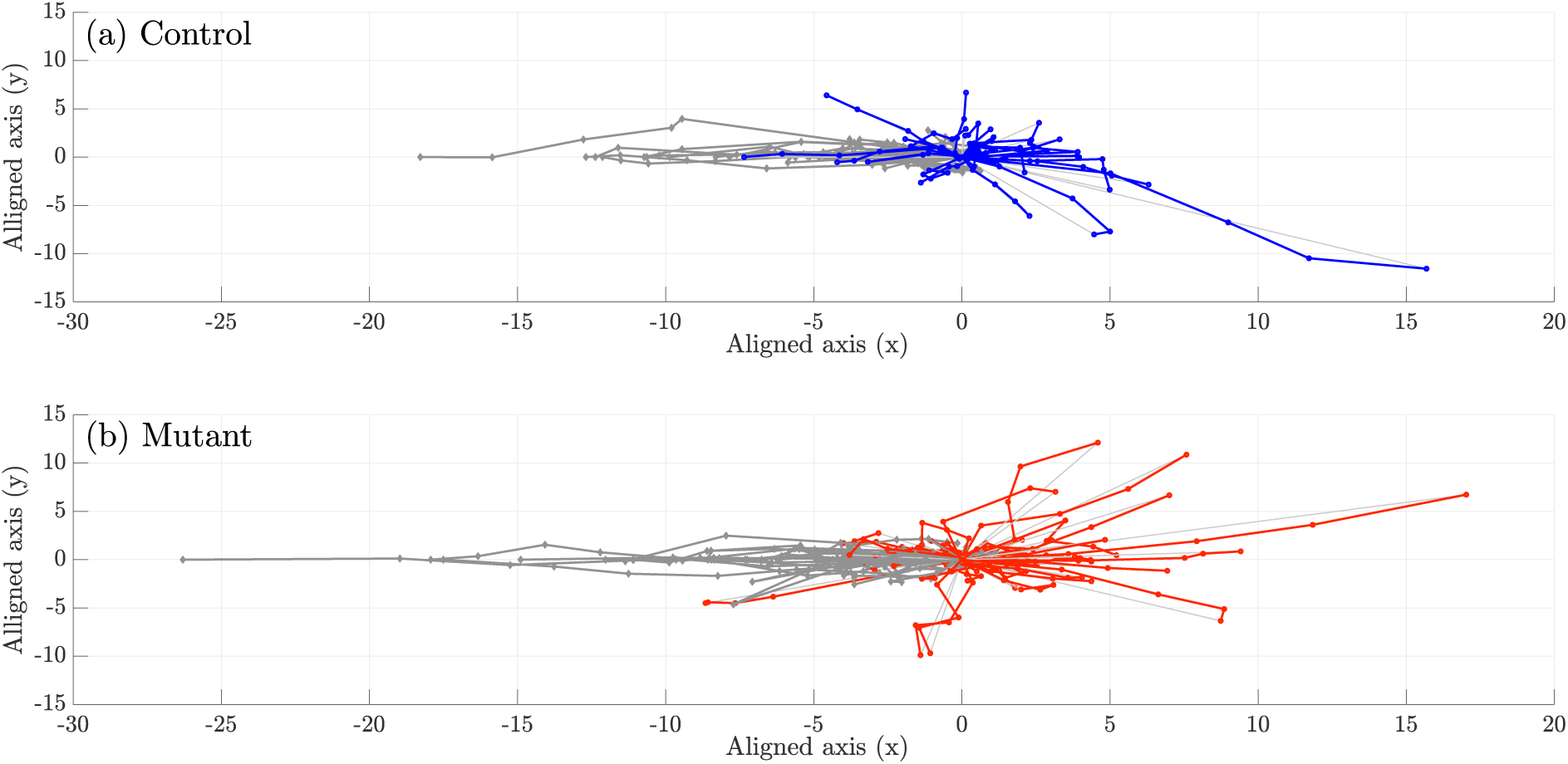
Comparison of aligned tracks for (a) Control and (b) Mutants interactions. Refer to figure 9 on how to read this figure.

Boxplots showing the change of direction angle *θ*_*x*_, time in clump *TC* and distances from the centre in the *x*′ are presented in figure 11. Notice that in figure 11(a), the angle *θ*_*x*_ for mutant interactions appear to be distributed with more cases towards the lower angles, or a smaller change of direction after the contact. However, on its own, this measurement did not provide a statistically significant difference. The data points where *θ*_*x*_ < 90 were chosen, as seen in figure 12. A t-test was calculated between the remaining angles showing statistical significance (*p* = 0.03 < 0.05). Finally, Figure 13 was created to visualise such difference in motion in the lower angles, *θ*_*x*_ < 90, but representing the distribution of angles in polar coordinates.

**Figure 11:**
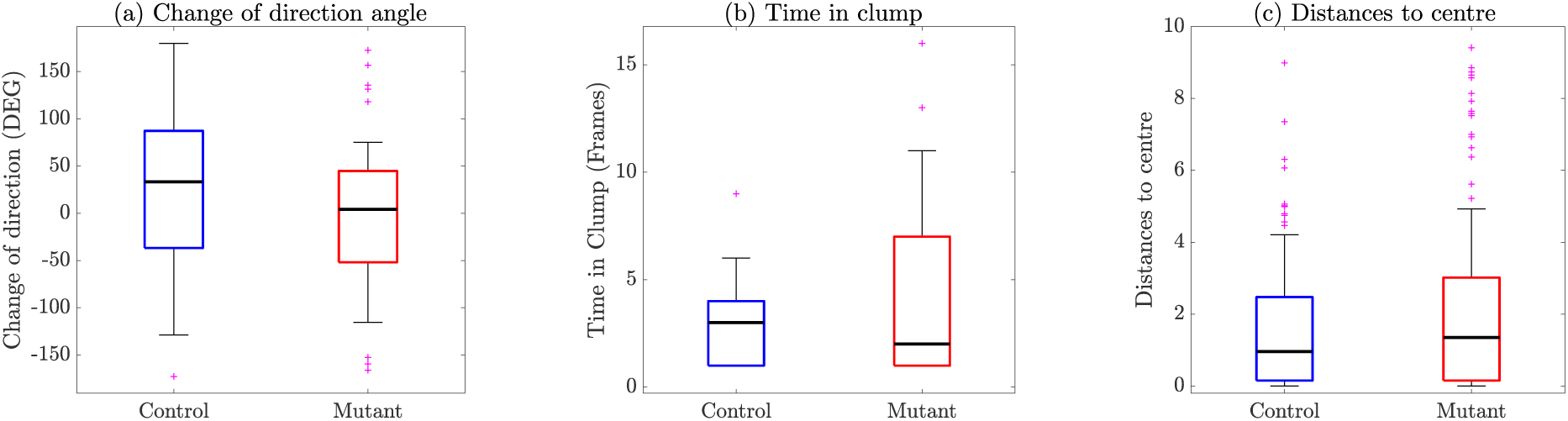
Comparison of relevant variables between Control (blue) and Mutant (red) interactions. (a) Change of direction angle, *θ*_*x*_, coming from figure 10. (b) Time in clump *TC* in frames. Finally, (c) shows the distances to the centre.

**Figure 12:**
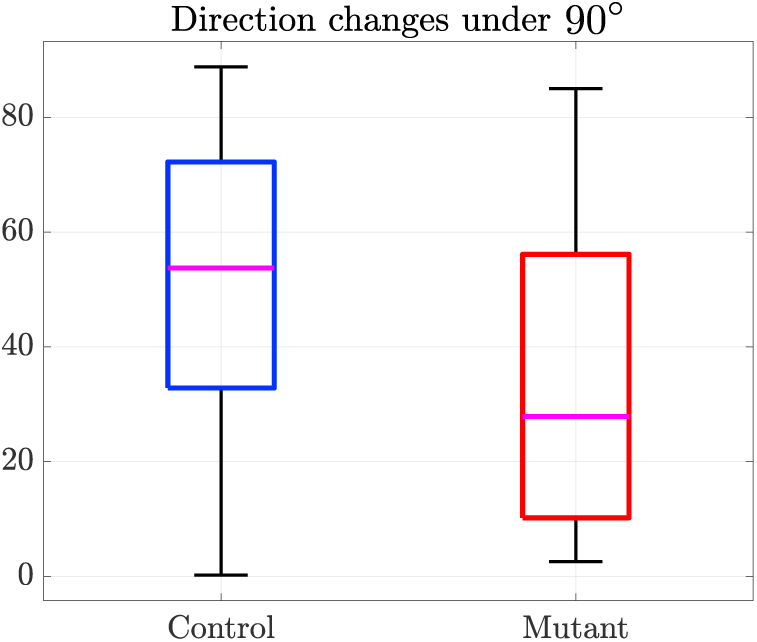
Change of direction differences between Control (blue) and Mutant (red) interactions for angles under 90°. After observation of figure 10(a), the smaller angles show a significant difference between the control and mutant interactions.

**Figure 13:**
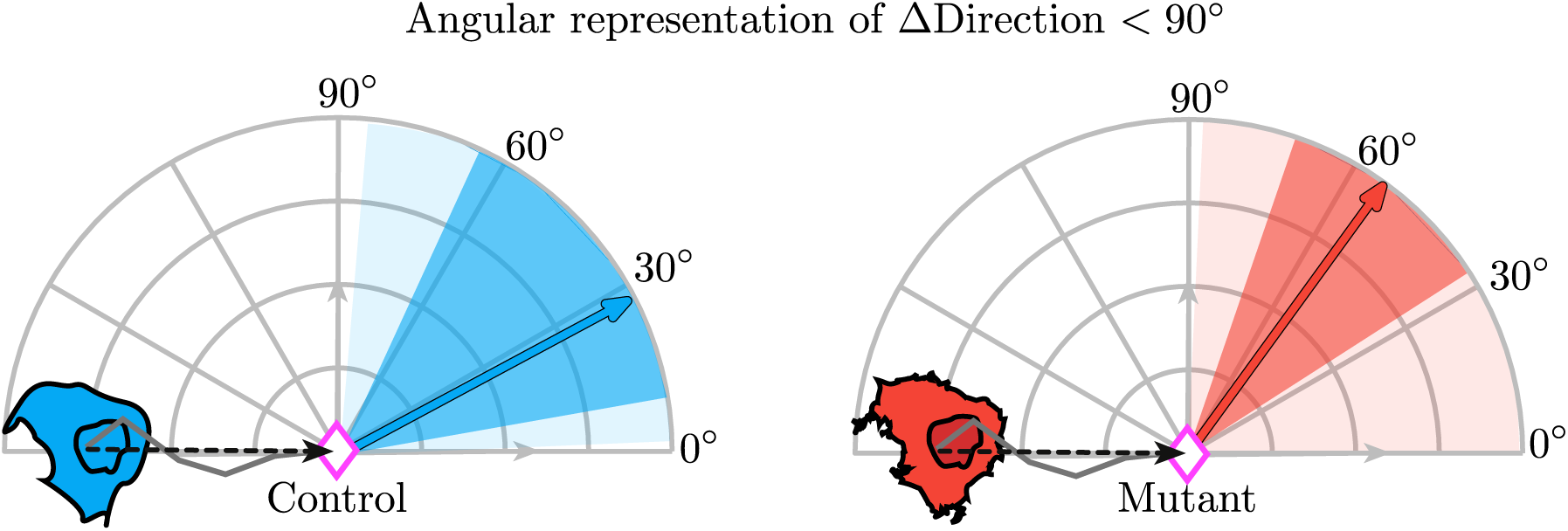
Angular representation of the change of direction for angles under 90°. These plots intend to represent the distributions seen in the boxplots of figure 12, but in a representation similar to that of Figure 10. The distribution of angles is represented by different shades of blue (control) or red (mutant), where the lighter shade represents the whiskers in a boxplot, the darker shade represents the box and the arrow represents the median.

## 5 DISCUSSION

This work presents a comparison of the movement that follows a contact between two cells. The populations of cells that were observed were mutant and wild-type macrophages observed with fluorescent microscopy during migration of macrophages in *Drosophila* embryos. Observation of such datasets indicate that the number of interactions found per dataset is not always consistent. In many cases, problems with the segmentation of the fine microtubule arm-like structures described in [31] can be lost due to the postprocessing stages of the segmentation. In particular with these datsets, the focus would vary extensively (Figure 2) complicating part of the analysis. Whilst the number of interactions that were selected from the datasets is small, there was an indication that there could be differences between the mutant and the wild-type cells in the sense that the wild-type cells show a greater change of direction after interaction than the mutants. However, to obtain this result it was necessary to select only interactions under specific conditions as seen in figure 12. In Figure 13, the distributions of angles are clearly visible, showcasing the differences reported by the statistical test.

## 6 CONCLUSIONS

This work presents a use case of the software `macrosight`, allowing an objective comparison of the movement following a contact between two different cell populations. While encouraging results were found, the differences between cell populations were oncly conclusive in very specific conditions. Future work will concentrate in increasing the number of datasets, which will in turn increase the number of interactions. Additionally, a larger number of variables collected from the tracking should be explored, and the segmentation could be enhanced with a step detecting discrete alignment of microtubules, therefore increasing the accuracy of interactions detected.

## Acknowledgements

This work was funded by a Doctoral Studentship granted by the School of Mathematics, Computer Science and Engineering at City, University of London.

## Author Contributions

JASL and CCRA conceived and designed the experiments; JASL performed the experiments; JASL and CCRA analysed the data; BS contributed the materials and motivation for this work; JASL wrote the paper and JASL, CCRA, BJSS and BS revised the paper.

## Conflicts of Interest

The authors declare no conflict of interest. The founding sponsors had no role in the design of the study; in the collection, analyses, or interpretation of data; in the writing of the manuscript, or in the decision to publish the results.

## Abbreviations

The following abbreviations are used in this manuscript:

ΔDirection: Change of direction
GFP: Green Fluorescent Probe
MIUA: Medical Image Understanding and Analysis
*𝒯*_*i*_: Track acquired with identifier *i*
*TC*: Time in clump

## Sample availability

Samples of the data and the code of the `macrosight` framework are available at https://github.com/alonsoJASL/macrosight under a GNU 3 open source license, or upon request to the corresponding author.

